# Faecal virome of the Australian grey-headed flying fox from urban/suburban environments contains novel coronaviruses, retroviruses and sapoviruses

**DOI:** 10.1101/2022.07.06.498921

**Authors:** Kate Van Brussel, Jackie E. Mahar, Ayda Susana Ortiz-Baez, Maura Carrai, Derek Spielman, Wayne S. J. Boardman, Michelle L. Baker, Julia A. Beatty, Jemma L. Geoghegan, Vanessa R. Barrs, Edward C. Holmes

**Affiliations:** Sydney Institute for Infectious Diseases, School of Life & Environmental Sciences and School of Medical Sciences, The University of Sydney, NSW, 2006, Australia; Jockey Club College of Veterinary Medicine & Life Sciences, City University of Hong Kong, Kowloon Tong, Hong Kong, SAR China; School of Veterinary Science, Faculty of Science, University of Sydney, Sydney, NSW 2006, Australia; School of Animal and Veterinary Sciences, Faculty of Science, Engineering and Technology, University of Adelaide, Adelaide, SA 5371, Australia; CSIRO Australian Centre for Disease Preparedness, Health and Biosecurity Business Unit, Geelong, VIC 3220, Australia; Department of Microbiology and Immunology, University of Otago, Dunedin 9010, New Zealand; Institute of Environmental Science and Research, Wellington 5022, New Zealand; Centre for Animal Health and Welfare, City University of Hong Kong, Kowloon Tong, Hong Kong, China

**Keywords:** Coronavirus, sapovirus, retrovirus, faecal, mammalian, grey-headed flying fox

## Abstract

Bats are important reservoirs for viruses of public health and veterinary concern. Virus studies in Australian bats usually target the families *Paramyxoviridae, Coronaviridae* and *Rhabdoviridae*, with little known about their overall virome composition. We used metatranscriptomic sequencing to characterise the faecal virome of grey-headed flying foxes from three colonies in urban/suburban locations from two Australian states. We identified viruses from three mammalian-infecting (*Coronaviridae, Caliciviridae, Retroviridae*) and one possible mammalian-infecting (*Birnaviridae*) family. Of particular interest were a novel bat betacoronavirus (subgenus *Nobecovirus*) and a novel bat sapovirus (*Caliciviridae*), the first identified in Australian bats, as well as a potentially exogenous retrovirus. The novel betacoronavirus was detected in two sampling locations 1,375 km apart and falls in a viral lineage likely with a long association with bats. This study highlights the utility of unbiased sequencing of faecal samples for identifying novel viruses and revealing broad-scale patterns of virus ecology and evolution.

## 1. Introduction

Bats (order Chiroptera) are one of the largest mammalian orders with a unique physiology adapted for flight. The number of bat colonies in urban habitats has increased in recent decades, leading to more frequent interactions with humans, companion animals and livestock that have in turn facilitated outbreaks of zoonotic disease (Plowright et al., 2011). This process has been dramatically highlighted by the emergence of severe acute respiratory syndrome coronavirus 2 (SARS-CoV-2) and the detection of SARS-like coronaviruses in Asian bat populations (Temmam et al., 2022, Zhou et al., 2021, Zhou et al., 2020, Wacharapluesadee et al., 2021, Murakami et al., 2020). In addition, bats have been associated with the emergence of Hendra virus (Halpin et al., 2000), Nipah virus (Yob et al., 2001), lyssaviruses (Botvinkin et al., 2003, Gould et al., 1998) and SARS-CoV (Li et al., 2005). In turn, these outbreaks have led to increased sampling of bat species, and the widespread use of metagenomic sequencing has enabled more detailed exploration of the bat virome (Wu et al., 2016, Hardmeier et al., 2021, Van Brussel and Holmes, 2022).

In Australia, bat species of the genus *Pteropus* are reservoir hosts for Hendra virus and Menangle virus, zoonotic pathogens of the family *Paramyxoviridae* (Halpin et al., 2000, Philbey et al., 1998), as well as Australian bat lyssavirus, a zoonotic virus of the *Rhabdoviridae* that causes rabies in mammals (Gould et al., 1998). Studies of viruses in bats in Australia have largely focused on these virus families and recently identified a new member of the *Paramyxoviridae* – Cedar virus – as well as a novel genotype of Hendra virus (Wang et al., 2021, Marsh et al., 2012). Although important, these studies lack information on overall virome composition, particularly those virus families not included in targeted PCR studies.

The grey-headed flying fox (*Pteropus poliocephalus*), a member of the megabat family Pteropodidae and native to Australia, is a species of importance in the context of zoonotic viruses. Grey-headed flying foxes are distributed throughout the eastern coastline of Australia (Queensland, New South Wales and Victoria) and more recently a colony was established in Adelaide (South Australia). Grey-headed flying foxes feed on fruit, pollen and nectar and roost in large colonies, sometimes sharing roosting locations with other species of *Pteropus*, allowing intraspecies and interspecies virus transmission (Timmiss et al., 2021). Roosting sites are commonly located alongside human communities including in densely populated urban settings (Williams et al., 2006). As numerous viruses are transmitted by faeces and other excretions, the co-habitation between bats and humans likely increases the risk of zoonotic spill-over.

Herein, we used metatranscriptomic sequencing of faecal samples to describe the community of viruses present in the gastrointestinal tract of grey-headed flying foxes from three sampling locations in two Australian states – Centennial Park and Gordon in Sydney, New South Wales, and the Botanic Park, Adelaide in South Australia. Specifically, to reveal the composition and abundance of viruses in bats residing in metropolitan areas we sampled roosting sites either located in a residential setting or in parks that are frequented by humans.

## 2. Methods

### Sample collection

Faecal samples were collected from grey-headed flying fox roosting sites in three regions of Australia: Centennial Parklands, Centennial Park New South Wales (NSW), Gordon NSW, and Botanic Park, Adelaide parklands, Adelaide, South Australia (Table 1, Fig. 1A). Sampling was conducted over two dates in 2019 for the Centennial Park and Gordon sites, while the roosting site in the Adelaide parklands was sampled over several months in 2019 (Table 1). A plastic sheet of approximately 3 × 5 metres was placed under densely populated trees the night before collection. The following morning samples captured by the plastic sheet were placed into 2 mL tubes and immediately stored at -80°C until processing. Any faecal sample touching or submerged in urine was discarded.

**Table 1.** Sampling overview, including number of samples allocated to sequencing pools and sequencing metadata.

| Location | Sampling date | Pool no. | No. of samples | No. of reads | No. of contigs |
| --- | --- | --- | --- | --- | --- |
| **Centennial Park, NSW**<br>33. 89999°S,<br>151.23592°E | 5 February 2019 | 01 | 12 | 24,732,494 | 159,527 |
|  |  | 02 | 9 | 35,835,953 | 147,425 |
|  |  | 03 | 9 | 31,960,624 | 107,431 |
|  | 26 February 2019 | 04 | 9 | 19,833,973 | 111,196 |
|  |  | 05 | 11 | 31,410,836 | 136,180 |
|  |  | 06 | 9 | 29,318,213 | 105,118 |
|  |  | 07 | 10 | 19,160,704 | 90,339 |
| **Gordon, NSW**<br>33.75065°S,<br>151.16242°E | 12 March 2019 | 01 | 12 | 52,605,108 | 89,247 |
|  |  | 02 | 12 | 48,784,843 | 50,574 |
|  |  | 03 | 9 | 27,396,450 | 118,509 |
|  | 26 March 2019 | 04 | 11 | 36,591,148 | 181,524 |
|  |  | 05 | 12 | 36,815,461 | 146,466 |
|  |  | 06 | 12 | 52,934,611 | 97,013 |
|  |  | 07 | 10 | 37,980,832 | 156,960 |
| **Adelaide, SA**<br>34.91571°S,<br>138.6068°E | 2019 | 01 | 8 | 25,977,712 | 135,969 |
|  | 2019 | 02 | 9 | 21,113,731 | 113,546 |

**Fig. 1.**
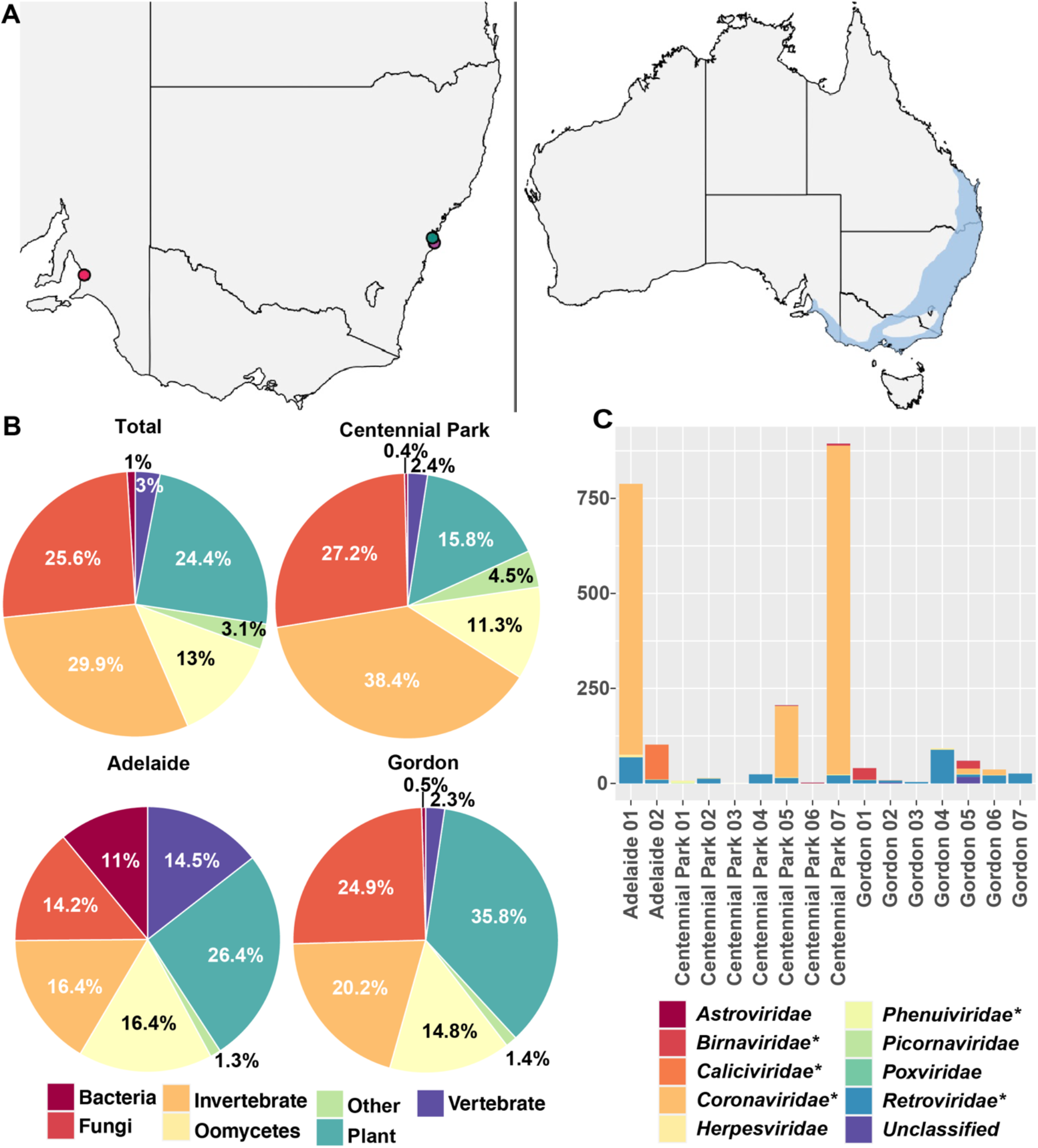
Overview of sampling sites and bat faecal sample composition. (A) Sampling locations in Australia (left) and distribution map of the grey-headed flying fox (right) (IUCN, 2021). (B) Likely hosts of viral contigs based on host designation of the closest relatives in the NCBI non-redundant protein database. (C) Read abundance presented as reads per million (RPM) for the vertebrate-associated virus sequences for each library and separated by virus family. The virus families discussed in this study are highlighted with an asterisk.

### RNA extraction, sequencing and read processing

Faecal samples were homogenised using the Omni Bead Ruptor 4 with 1.44 mm ceramic beads (Omni international). Total RNA was extracted from each sample individually using the RNeasy Plus Mini Kit (Qiagen) following the manufacturer’s protocol. RNA was pooled in equimolar ratios and separated by sampling location, date and RNA concentration (Table 1). Ribosomal RNA was depleted, and libraries constructed using the Illumina Stranded Total RNA Prep with Ribo-Zero Plus (Illumina) preparation kit. Libraries were sequenced as 150 bp paired-end on the Illumina Novaseq 6000 platform at the Australian Genome Research Facility (AGRF). Read ends with a quality score of below 25 phred and adapter sequences were removed using cutadapt v1.8.3 (Kechin et al., 2017). Sortmerna v4.3.3 was used to remove 5S and 5.8S, eukaryotic 18S and 23S, bacterial 16S and 23S, and Archaea 16S and 23S ribosomal RNA (rRNA) reads (Kopylova et al., 2012). The filtered reads were then *de novo* assembled using Megahit v1.1.3 (Li et al., 2015) and contigs were compared to the non-redundant protein database using diamond v2.0.9. The Genemark heuristic approach (Besemer and Borodovsky, 1999, Zhu et al., 2010) and information from closely related viruses were used to predict genes and annotate genomes. Intact retrovirus genomes were detected using an in-house pipeline (Chang et al., manuscript in preparation). The Geneious assembler (Geneious Prime version 2022.1.1) was used to reassemble megahit contigs from multiple libraries for bat faecal associated retrovirus 2 (see Results). The final sequence for bat faecal associated retrovirus 2 (see Results) was determined by mapping reads from all libraries to the reassembled genome on Geneious Prime using a 0% (majority) threshold for the final consensus sequence.

### Abundance estimation

Virus and host abundance were estimated by mapping non-rRNA reads from each library to assembled contigs, and to the COX1 gene (accession no. KF726143) from the *P. alecto* (Black flying fox) genome using Bowtie2 v2.3.4.3 (Langmead and Salzberg, 2012). The impact of index-hopping was minimised by excluding the read abundance count for a contig in any library that was less than 0.1% of the highest read count for that assembled contig in any other library.

### Phylogenetic analysis

Virus amino acid sequences were aligned with related sequences (i.e., representing the same virus family and/or genus) retrieved from the NCBI/GenBank database using MAFTT v7.450 (Katoh and Standley, 2013) and the E-INS-I algorithm (Katoh et al., 2005). The partial RdRp sequence of P. alecto/Aus/SEQ/2009 was retrived from Smith et al. (2016). The gappyout method in TrimAL v1.4.1 was used to remove ambiguous regions in the alignment (Capella-Gutiérrez et al., 2009). Maximum likelihood trees of each data set were inferred using IQ-TREE v1.6.7 (Nguyen et al., 2014), employing the best-fit amino acid substitution model determined by the ModelFinder program (Kalyaanamoorthy et al., 2017) in IQ-TREE. Nodal support was accessed using 1000 ultrafast bootstrap replicates (Hoang et al., 2017). Any virus sequence with over 90% nucleotide similarity to another detected here was excluded from the phylogenetic analysis.

## 3. Results

### Virome overview

In total, 164 faecal samples allocated to 16 libraries underwent metatranscriptomic sequencing. This generated 19,160,704 to 52,934,611 reads per library (average of 33,278,293 reads) after read filtering (Table 1). Reads were *de novo* assembled into 50,574 to 181,524 contigs (average of 121,689 contigs) per library (Table 1). A total of 5,933 contigs were assigned as of viral origin across all the libraries. The samples collected at Centennial Park, Sydney produced the most viral contigs, with 3,216 identified from 65 virus families (Supplementary Fig. 1). The Gordon, NSW sample site produced 2,399 virus contigs from 66 virus families, while the Adelaide site contained 318 virus contigs from 33 virus families, although this site had only two sequencing libraries comprising 17 faecal samples, compared to seven sequencing libraries in each of the other two locations (69 faecal samples from Centennial Park, 78 from Gordon) (Table 1, Supplementary Fig. 1). Screening of the NCBI protein database revealed assembled virus contigs were mostly associated with infection of invertebrates (29.9% of total contigs), fungi (25.6%), plants (24.4%), and oomycetes (13%), representing 79 virus families (Fig. 1B, Supplementary Fig. 1). These viruses were most likely associated with host diet and differed in frequency depending on sampling site (Fig. 1B, Supplementary Fig. 1). The plant, fungal, and oomycete-associated viruses, as well as those likely to be bacteriophage (including the picobirnaviruses) were not considered further. Importantly, however, we also identified sequences from viruses likely associated with mammalian infection (3% overall), including near complete genomes from members of the *Coronaviridae, Caliciviridae* and *Retroviridae* (Fig. 1B).

### Mammalian-associated viruses

We detected contigs from nine viral families likely to infect mammals (Fig. 1C). The *Coronaviridae* and *Retroviridae* were particularly abundant and present in five and 13 libraries, respectively (Fig. 1C). Members of the *Birnaviridae* and *Caliciviridae* were also abundant in specific libraries (Fig. 1C). The remaining mammalian-associated viral families were only detected at low abundance and the contigs were not of sufficient length for further characterisation.

### Novel betacoronavirus (*Coronaviridae*)

A novel complete betacoronavirus genome (single-strand, positive-sense RNA virus; +ssRNA) – provisionally denoted bat faecal coronavirus CP07/aus/1 – was identified in a sequencing library sampled from Centennial Park (pool no. 07) and in a sequencing library from Adelaide (pool no. 01). These two sequences exhibited 99.8% identity over the complete viral genome indicating that they represent the same species. Additionally, three sequences with 99.2-100% sequence identity to CP07/aus/1 were identified in an additional Centennial Park library (pool no. 05).

CP07/aus/1 contains ten ORFs in the arrangement ORF1a, ORF1ab, spike, NS3, envelope, matrix, nucleocapsid, NS7a, NS7b and NS7c. Transcription Regulatory Sequences (TRS) preceeded all ORFs. Additional bat coronavirus contigs ranging from 318 to 1,309 bp were detected in sequencing libraries from two Gordon sampling locations. These short contigs shared 40-95% amino acid identity to CP07/aus/1. Three of these contigs contained RdRp or spike amino acid sequences of sufficient length for phylogenetic analysis, and these were provisionally denoted bat faecal coronavirus G05/aus/1, G05/aus/2 and G05/aus/3. Based on phylogenetic analysis of the RNA-dependent RNA polymerase (RdRp) and/or spike protein, the novel betacoronaviruses detected here fell within the *Betacoronavirus* subgenus *Nobecovirus* (Fig. 2) and were most closely related to P.alecto/Aus/SEQ/2009 (for which only a partial RdRp is available) sampled from a black flying fox in south east Queensland, Australia (Smith et al., 2016) and to Pteropus rufus nobecovirus sampled from a flying fox in Madagascar (accession no. OK067319; Fig. 2) (Kettenburg et al., 2022). CP07/aus/1 had 83% amino acid identity to Pteropus rufus nobecovirus over the complete ORF1ab replicase and 97% to P.alecto/Aus/SEQ/2009 over the partial RdRp. Amino acid identity to Pteropus rufus nobecovirus over the spike and non-structural proteins was 72% and 58%, respectively. The RdRp of G05/aus/1 shared 95% amino acid identity to CP07/aus/1, while the partial spike proteins of G05/aus/2 and G05/aus/3 shared 57% and 63% amino acid identity to CP07/aus/1, respectively. It is possible that G05/aus/1 and G05/aus/2 represent transcripts from the same virus, while G05/aus/3 represents a different species to CP07/aus/1. However, this could not be confirmed as the G05/aus/3 genome was incomplete. Regardless, it is clear from the spike protein phylogeny that at least three different coronaviruses are circulating in the bats sampled here.

**Fig. 2.**
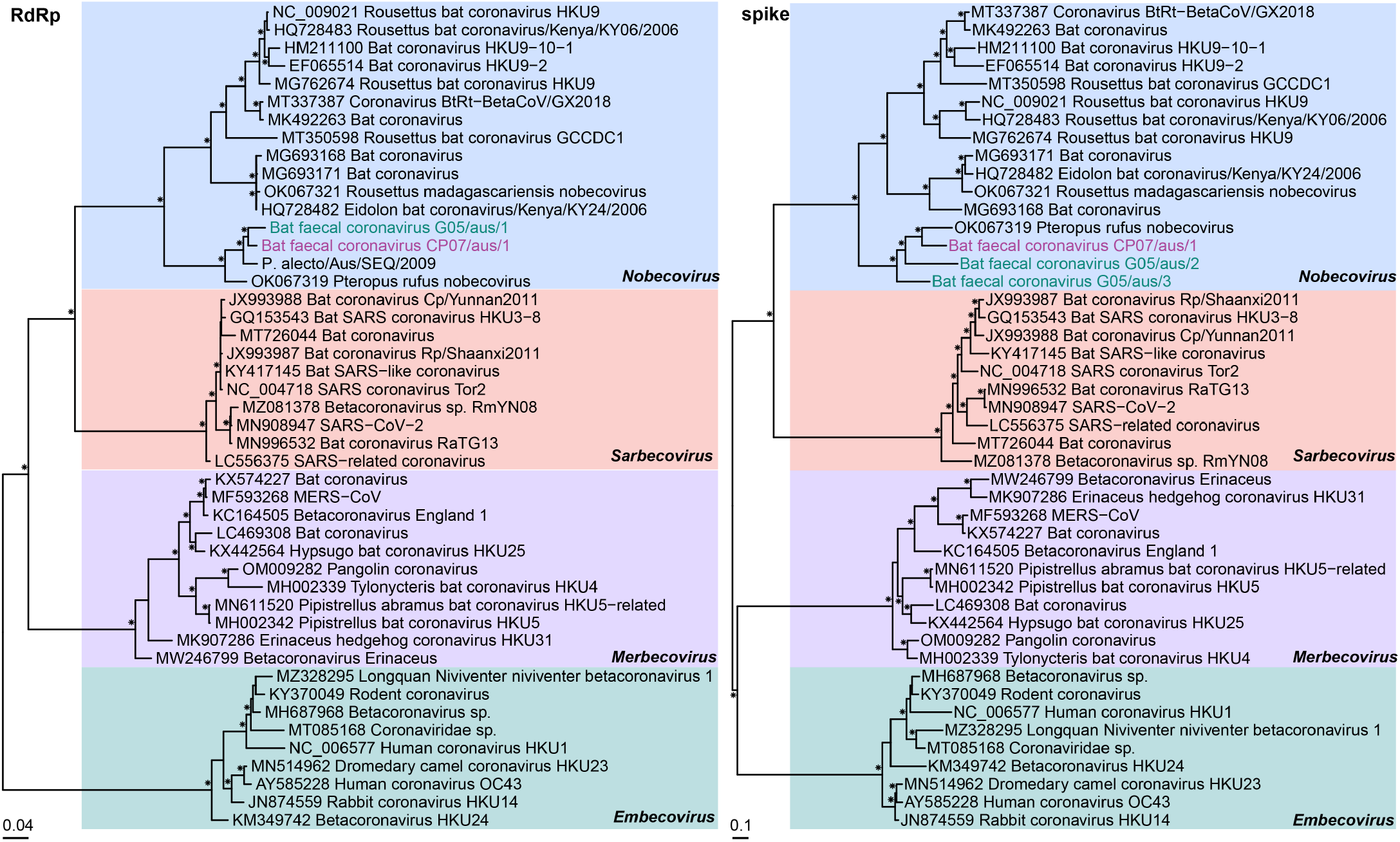
Phylogenetic relationships of the novel bat betacoronaviruses based on the amino acid sequences of the RdRp and spike protein. Amino acid alignment lengths were 832 and 1,092 residues for the RdRp and spike protein, respectively. Representative betacoronavirus sequences from this study are coloured by sampling location (Centennial Park, Sydney – purple, and Gordon – green) and the subgenera are highlighted. Bootstrap values >70% are represented by the symbol shown at the branch node. The tree is rooted at midpoint for clarity and the scale bar represents the amino acid substitutions per site.

### Novel sapovirus (*Caliciviridae*)

A near complete genome of a novel sapovirus (*Caliciviridae*, +ssRNA virus), tentatively named bat faecal sapovirus Ad02/aus/1, was detected in a sequencing library sampled from Adelaide (pool no. 2). Nine additional bat sapovirus sequences ranging from 340 to 783 bp were detected in the same sequencing library. The nine sequences shared 66-74% nucleotide and 76-81% amino acid identity to Ad02/aus/1 over the polyprotein, suggesting the presence of additional diverse sapoviruses. The near complete Ad02/aus/1 genome is 7,254 bp and contains two ORFs encoding a polyprotein (near complete with likely 45 residues missing from the 5’ end), and the VP2. Ad02/aus/1 exhibited 44.8% amino acid identity in the partial polyprotein to its closest relative – Bat sapovirus Bat-SaV/Limbe65/CAM/2014 (accession no. KX759620) – detected in the faeces of *Eidolon helvum* bats in Cameroon, Africa (Yinda et al., 2017). Phylogenetic analysis of the RdRp and VP1 revealed a clustering of bat sapoviruses in both trees that included the novel Australian bat sapoviruses found here (Fig. 3). Bat sapoviruses have been assigned to the putative genogroups GXIV, GXVI, GXVII, GXVIII and GXIX based on VP1 phylogeny and amino acid sequence identities. Using the same criteria, the novel sapovirus Ad02/aus/1 identified here should be assigned to its own genogroup, putatively named GXX, which would also include the partial VP1 Ad02/aus/4 sequence (Supplementary Table 1, Fig. 3).

**Fig. 3.**
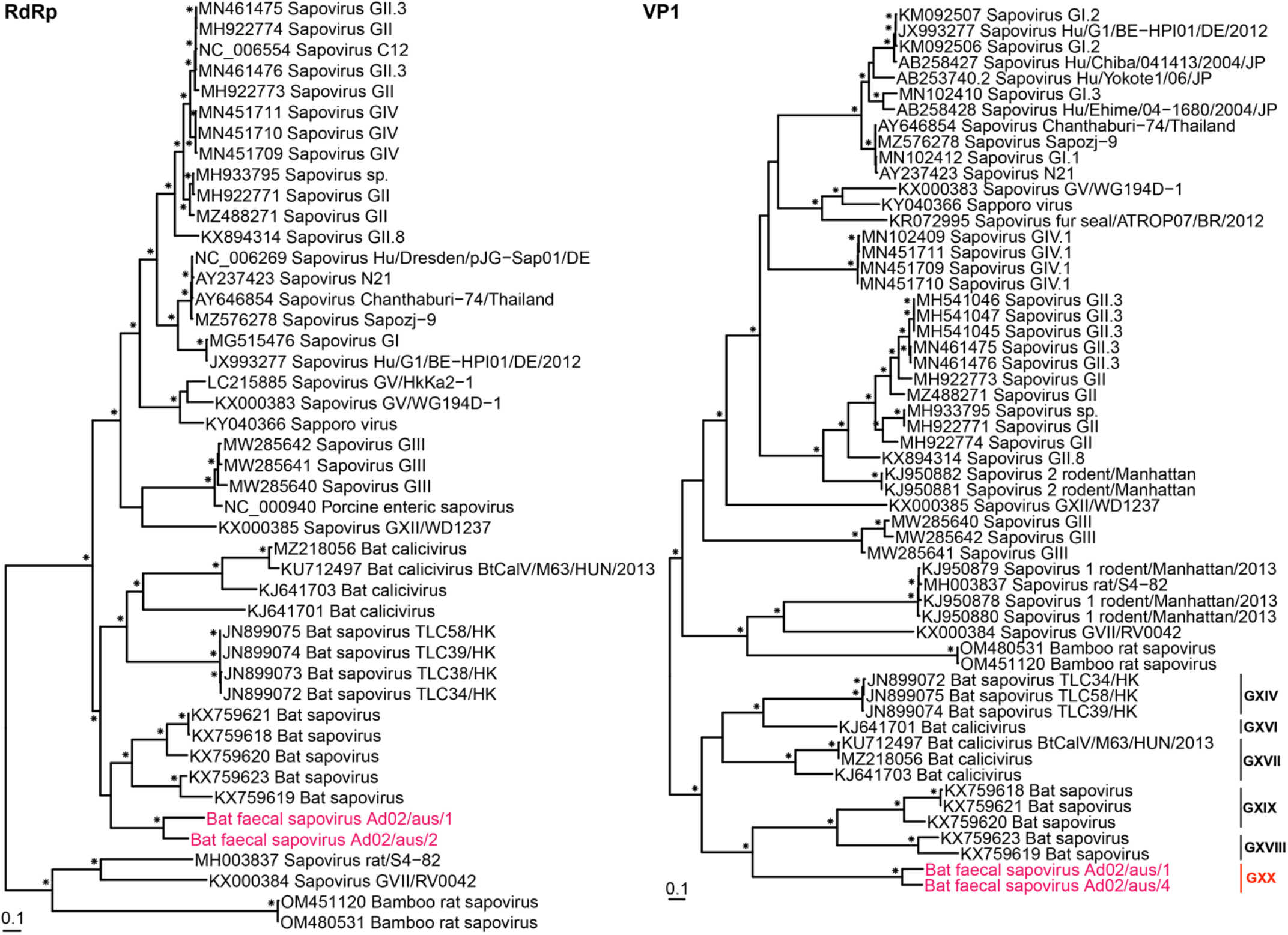
Phylogenetic relationships of the novel bat sapoviruses using the amino acid sequences of the RdRp and VP1. Amino acid alignment lengths were 491 and 623 residues for the RdRp and VP1, respectively. Bat sapoviruses from this study are coloured by sampling location (Adelaide – pink) and bootstrap values >70% are represented by the symbol shown at the branch node. The putative bat sapovirus genogroups are displayed to the right of the VP1 tree and our proposed putative genogroup is coloured in red. The trees are rooted at midpoint for clarity and the scale bar represents the amino acid substitutions per site.

### Novel birna-like virus (*Birnaviridae*)

Sequences related to the *Birnaviridae* (double-stranded RNA viruses; dsRNA) were detected in one Centennial Park and two Gordon libraries. All the birna-like virus sequences identified in the Centennial Park and Gordon libraries shared >99% nucleotide identity, and the complete coding region of segment B, which encodes the RdRp, was obtained from one library (Gordon 05). The *Birnaviridae* segment A that encodes the polyprotein and a small overlapping ORF was not identified in our data. Phylogenetic analysis revealed that the birna-like virus RdRp sequence, denoted G05/aus/1, was most closely related (50% amino acid identity) to the disease-causing virus Chicken proventricular necrosis virus (Fig. 4) (Guy et al., 2011), forming a distinct clade that is distantly related to the birnaviruses that infect a wide range of hosts.

**Fig. 4.**
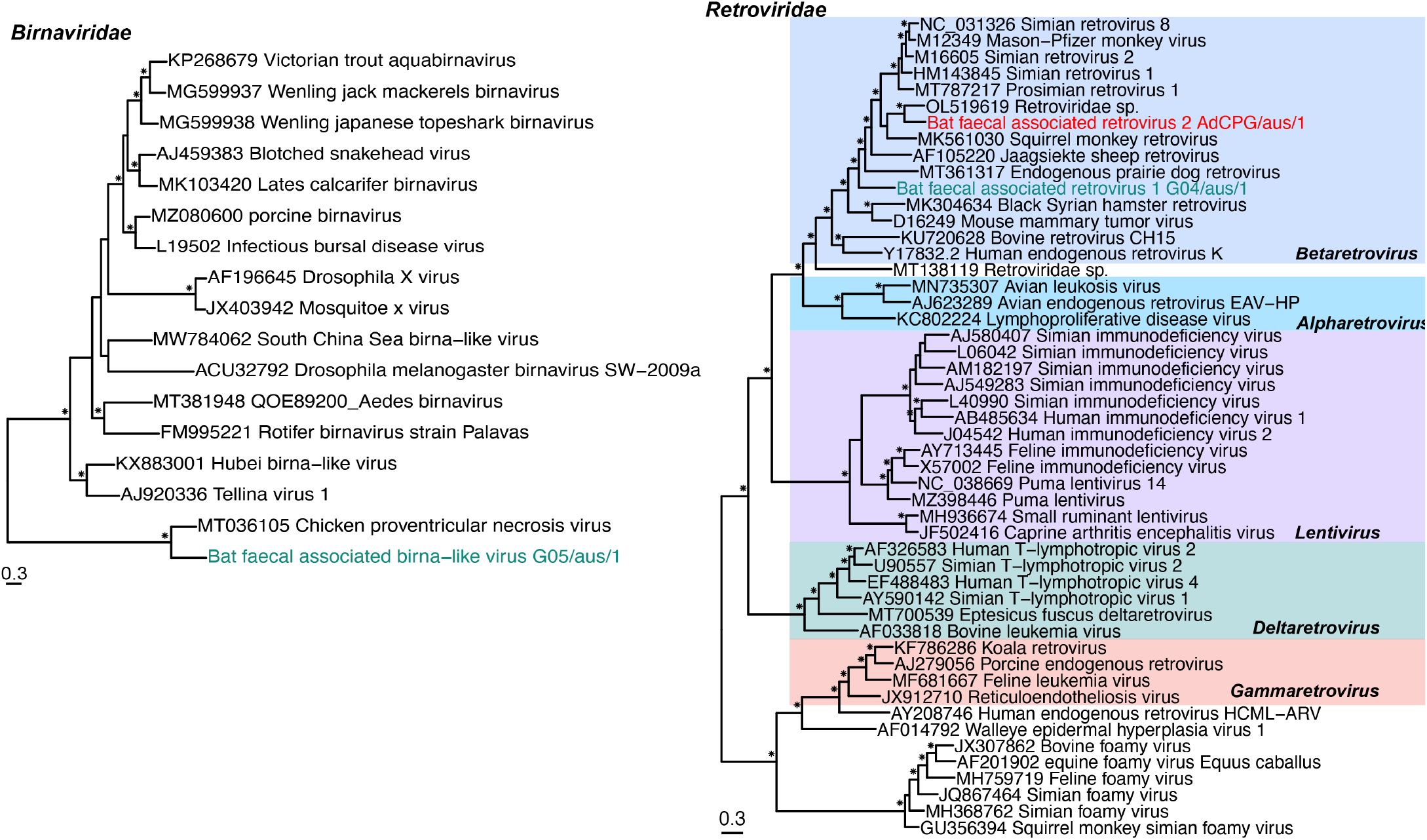
Phylogenetic analysis of the birna-like virus and bat retroviruses based on the RdRp and pol amino acid sequences, respectively. The *Birnaviridae* RdRp sequence alignment was 767 amino acid resides in length while the *Retroviridae* pol alignment comprised 1,356 residues. The viruses from this study are coloured by sampling location (Gordon – green) and the reassembled retrovirus sequence is in red (to indicate multiple locations). The *Retroviridae* genera are highlighted and bootstrap values >70% are represented by the symbol shown at the branch node. The tree is midpoint rooted for clarity, with the scale bar representing the amino acid substitutions per site.

### Bat retrovirus (*Retroviridae)*

A near complete genome of a retrovirus was identified in Gordon library 04 and provisionally named bat faecal associated retrovirus 1 G04/aus/1. Four ORFs were observed over the 7,455 bp genome and assigned as the gag, pro, pol and env genes based on the presence of conserved domains. In the pro gene we were able to identify an active site motif DTGAD predominately observed in functional retroviruses, and a helix motif GRDVL (Turnbull and Douville, 2018). We were unable to identify complete long terminal repeat (LTR) regions in the 7,455 bp genome, although this may be due to incomplete assembly at the 5’ and/or 3’ end, rather than a true absence of LTRs. Importantly, as the four ORFs contained the appropriate retrovirus conserved domains and were uninterrupted by stop codons, it is possible that G04/aus/1 is potentially exogenous and functional. A BLASTn analysis of the complete G04/aus/1 genome revealed no match to any bat reference genome on NCBI/GenBank. G04/aus/1 exhibited 56% amino acid identity in the pol protein to its closest relative, Simian retrovirus 2 (accession M16605), a presumably exogenous retrovirus (Thayer et al., 1987). The abundance for this novel retrovirus in the Gordon 04 library was 67 RPM (2,457 reads) (Fig. 1C).

A further near complete retroviral genome was identified by reassembling 31 partial contig sequences from 12 libraries from all three sample locations. This Bat faecal associated retrovirus 2 AdCPG/aus/1 is 6,630 bp and contains four open reading frames encoding the gag, pro, pol and env genes. It also contains the conserved domains expected in functional retroviruses, although the terminal end of the env gene is missing (either from true truncation or incomplete assembly). The virus is most closely related to AdCPG/aus/1 sampled from the lung tissue of Malayan pangolins (Ning et al., 2022). BLASTn analysis of the complete genome of AdCPG/aus/1 showed the absence of this genome in any bat reference genome on NCBI/GenBank. AdCPG/aus/1 reads were detected in 13 libraries (two Adelaide, four Centennial Park and seven Gordon) and the abundance in each library ranged from 3.7 – 68.8 RPM (127 – 1786 reads) (Fig. 1C). Phylogenetic analysis of the pol protein that contains the reverse transcriptase (RT) domain revealed that G04/aus/1 and AdCPG/aus/1 fell within the genus *Betaretrovirus*, clustering with both exogenous and endogenous retroviruses associated with various mammalian species (Fig. 4).

### Invertebrate-associated viruses

We detected likely invertebrate-associated virus sequences from seven single-strand negative-sense RNA viruses (-ssRNA), three +ssRNA virus and one dsRNA virus families, in addition to the order *Bunyavirales* (-ssRNA). The virus sequences from the *Chuviridae, Lispiviridae, Artoviridae, Nyamiviridae, Xinmoviridae, Qinviridae, Disctroviridae* and *Iflaviridae* are not discussed further, although information on positive libraries is provided (Supplementary Fig. 1) and phylogenetic analysis was performed (Supplementary Fig. 2). Virus sequences from the *Orthomyxoviridae, Nodaviridae, Reoviridae* and *Bunyavirales* are considered further as these viral groups include mammalian-infecting viruses, are important vector-borne viruses, or are able to infect mammals experimentally (*Nodaviridae*, genus *Alphanodavirus*).

Orthomyxovirus (-ssRNA virus) segments were identified in five libraries from Centennial Park. Full coding regions for two polymerase segments – PB2 and PA – and the hemagglutinin segment 2 and nucleocapsid segment 5 were present in all libraries, although a full coding region for polymerase segment PB1 was only present in a single Centennial Park library. The three polymerase proteins of Centennial Park library 06 were used for phylogenetic analysis, which revealed that this sequence was most closely related to an orthomyxovirus sampled from jumping plant lice in Australia (Fig. 5) (Käfer et al., 2019). Nodaviruses (+ssRNA virus) were detected in five Centennial Park libraries and three Gordon libraries. Both the RNA1 (RdRp) and RNA2 segments were identified, including two sequences with the complete RdRp. Nodavirus CP01/aus/1 and CP02/aus/1 were related to a nodavirus sampled from birds in China (Zhu et al., 2022) and most likely belong to the same viral species, although these fragments were only 476 and 232 amino acids, respectively. The nodavirus CP07/aus/1 RdRp segment was related to a nodavirus from arthropod hosts from China (Fig. 5) (Shi et al., 2016). Gene segments related to the *Reoviridae* (dsRNA) were present in all Centennial Park, three Gordon and one Adelaide library. The reovirus VP1 Pol segments detected here were related, albeit distantly (∼40% amino acid identity) to reoviruses associated with ticks (Harvey et al., 2019, Vanmechelen et al., 2021), moths (Graham et al., 2006), bat flies (Xu et al., 2022) and the Asian citrus psyllid (Nouri et al., 2015) (Fig. 5).

**Fig. 5.**
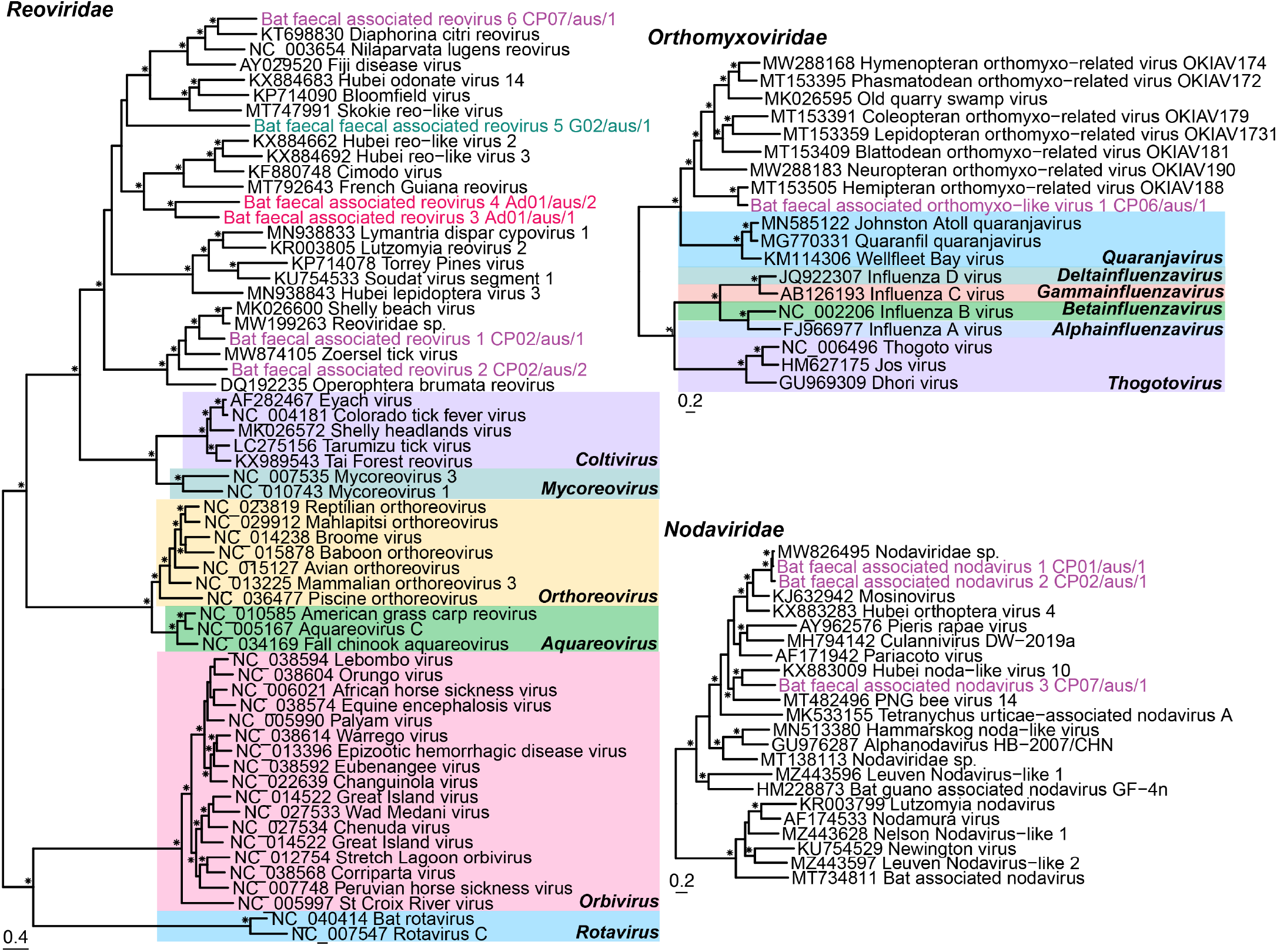
Phylogenetic analysis of the invertebrate-associated reoviruses, orthomyxoviruses and nodaviruses based on the VP1 Pol, concatenated PB2-PB1-PA and RdRp amino acid sequences, respectively. Amino acid alignment length were 1,020 residues for *Reoviridae*, 2,233 residues for the *Orthomyxoviridae* and 774 residues for the *Nodaviridae*. Viruses from this study are coloured by sampling location (Adelaide – pink, Centennial Park – purple and Gordon – green) and genera are highlighted in the *Reoviridae* and *Orthomyxoviridae* tress. Bootstrap values >70% are represented by the symbol shown at the branch node. The tree is rooted at midpoint for clarity and the scale bar represents the amino acid substitutions per site.

Finally, bunyavirus fragments were detected in all the Adelaide and Centennial Park libraries and six Gordon libraries. Eleven RdRp coding regions were used for phylogenetic analysis which revealed that two bunyavirus sequences fell into the *Phenuiviridae* and four were basal to that family, while two sequences fell into the *Phasmaviridae*, two were basal to the *Arenaviridae*, and one was basal to a grouping of five families (Fig. 6). The Adelaide bunyavirus Ad02/aus/1 was related to the plant associated genus *Tenuivirus* and the remaining 10 were related to invertebrate hosts (Fig. 6).

**Fig. 6.**
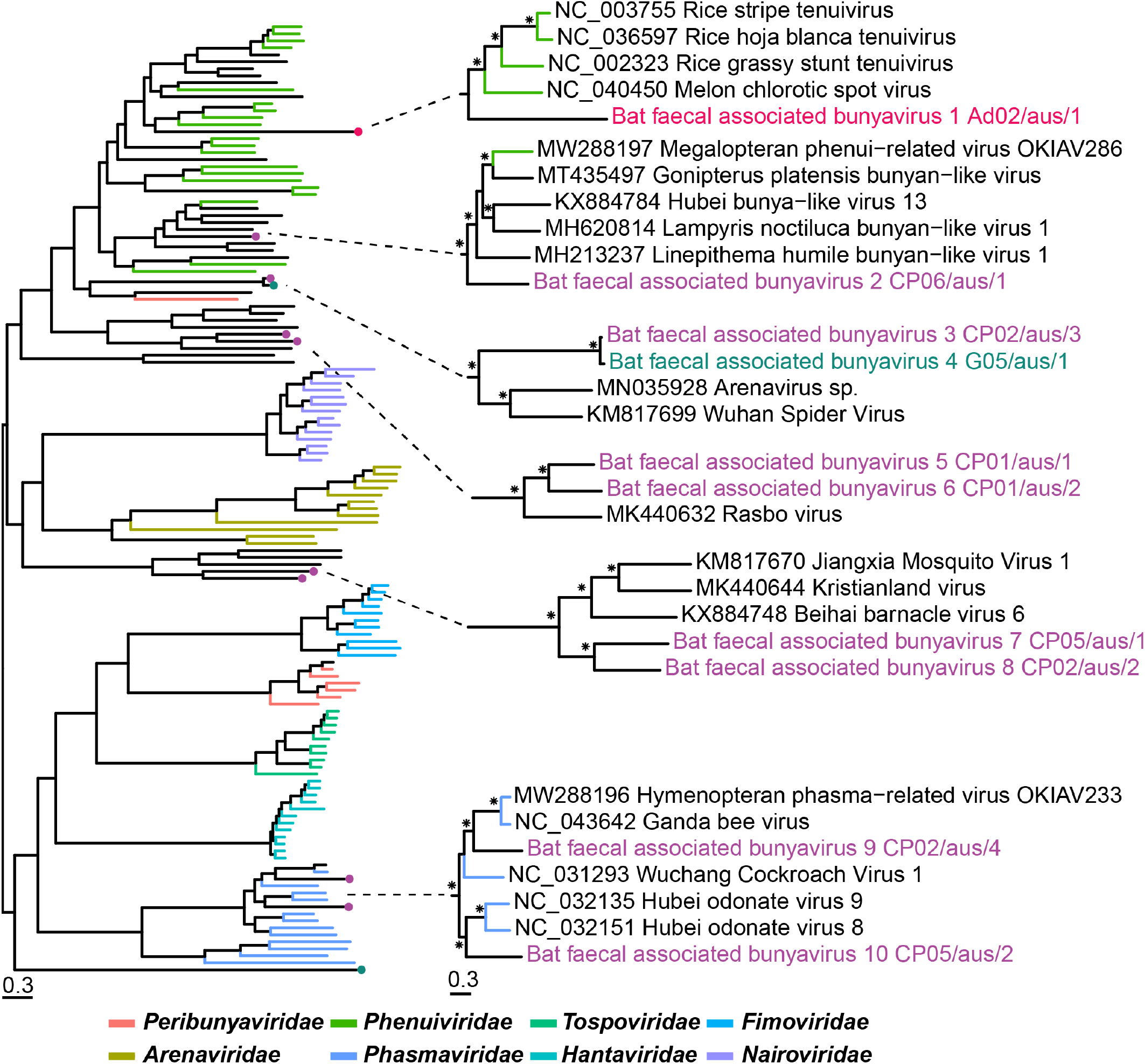
Phylogenetic analysis of viruses from the order *Bunyavirales*. The RdRp amino acid sequence was used to estimate phylogenetic trees and the alignment length was 1,434 amino acid residues. Viruses from this study are coloured by sampling location and bootstrap values >70% are represented by the symbol shown at the branch node. The tree is midpoint rooted for clarity and the scale bar represents the amino acid substitutions per site.

## 4. Discussion

Virological surveillance of bats in Australia has largely focused on screening for known zoonotic viruses such as Hendra virus and Australian bat lyssavirus, although the paramyxovirus Tioman virus, for which flying foxes are the natural host, and coronaviruses are also targeted (Boardman et al., 2020, Prada et al., 2019a, Smith et al., 2016). The primary aim of these studies is to identify specific viruses using either PCR or serological data. Although such surveillance has been successful in determining the active circulation of these specific viruses, these approaches necessarily have restricted capacity to detect novel or unexpected viruses, thus providing a limited understanding of viruses circulating in Australian bats. As bats are frequently found near human populations, they are of particular concern regarding potential zoonoses (Plowright et al., 2011, Williams et al., 2006, Halpin et al., 2000). Herein, we used metatranscriptomics to reveal the natural faecal virome of the grey-headed flying fox. Although most of the viruses identified were likely associated with bat diet, as expected from faecal sampling, we also identified viruses from three mammalian-associated families (*Coronaviridae, Caliciviridae, Retroviridae*) and one virus from the *Birnaviridae* family that may also have a mammalian association.

Both alpha- and betacoronaviruses have been identified in a variety of bat species (Smith et al., 2016, Prada et al., 2019b). Here, we characterised the complete genome of a betacoronavirus in grey-headed flying foxes that was closely related to two other betacoronaviruses sampled in flying foxes in Australia and Madagascar (Smith et al., 2016, Kettenburg et al., 2022). The current ICTV classification for coronavirus species states that less than 90% amino acid identity in the ORF1ab conserved replicase domains constitutes a new species. Although bat faecal coronavirus CP07/aus/1 shares high sequence similarity to another reported bat betacoronavirus, the P.alecto/Aus/SEQ/2009 sequence is only 146 amino acids in length, does not span the complete RdRp and is therefore difficult to classify. Accordingly, we suggest that betacoronavirus bat faecal coronavirus CP07/aus/1 represents a novel species, to which P.alecto/Aus/SEQ/2009 may also belong. The complete genome of this virus was found in both Adelaide and New South Wales (99.8% nucleotide similarity between the two genomes) and abundance counts were high in both locations (Fig. 1C), indicative of virus exchange between bat populations. Flying foxes are known to travel long distances to feed, roosting sites change depending on season, and in Australia several flying fox species share roosting sites (Timmiss et al., 2021), all of which provide opportunities for viruses to infect new individuals. Importantly, while we were only able to assemble the complete genome of one novel coronavirus, we identified partial genome fragments of at least two more diverse coronaviruses (Fig. 2), indicating that Australian bats carry a high diversity of coronaviruses as has been seen in other bat species.

This is the first report of a sapovirus in Australian bats. Previously, bat sapoviruses have been sampled from *Eidolon helvum* (Straw-coloured fruit bat) in Cameroon (Yinda et al., 2017) and Saudi Arabia (Mishra et al., 2019) and *Hipposideros Pomona* (Pomona leaf-nosed bat) from Hong Kong (Tse et al., 2012). Currently, the bat sapoviruses characterised have been from bats with no apparent disease (Tse et al., 2012, Yinda et al., 2017, Mishra et al., 2019). Whether this is the case here is unknown because the reliance on faecal sampling meant that there was no direct interaction with individual animals. The disease potential of bat sapoviruses should be investigated further as sapoviruses have been linked to acute gastroenteritis outbreaks in humans (Oka et al., 2015) and some animal sapoviruses are closely related to those found in humans (Mombo et al., 2014, Firth et al., 2014, Martella et al., 2008).

Until the metagenomic detection of porcine birnavirus (Yang et al., 2021) and porcupine birnavirus (He et al., 2022) it was believed the *Birnaviridae* infected fish, insects and birds exclusively (Crane et al., 2000, Da Costa et al., 2003, Chung et al., 1996, Brown and Skinner, 1996, Guy et al., 2011). We identified the segment B sequence of a novel bat faecal associated birna-like virus that was most closely related to a divergent pathogenic avian birnavirus (50% amino acid identity). Given its divergent phylogenetic position it is currently unclear whether this virus actively infects grey-headed flying foxes or is associated with a component of their diet or microbiome. While grey-headed flying foxes are not insectivores, the ingestion of insects through the consumption of fruit and nectar seems likely given the high number of invertebrate, plant and fungi viruses sequenced here (Fig. 1B, Supplementary Fig. 1). The moderate abundance values (30.5 and 19.9 RPM) cannot exclude either scenario as using a host reference gene such as COX1 for sequencing depth comparison may not be as reliable for faecal samples as it would be when analysing tissue. Further investigation is needed to determine the natural host of bat faecal associated birna-like virus and to determine what tissue types are affected.

Two intact, possibly exogenous retrovirus near complete genomes were also identified in this study and were most closely related to mammalian infecting retroviruses from the genus *Betaretrovirus*. Six retroviruses have been previously characterised from Australian bat brain tissue and excretions (including faeces), all from the genus *Gammaretrovirus* (Hayward et al., 2020, Cui et al., 2012) and hence highly divergent from the viruses identified here. Although the exogenous status needs to be confirmed, it is possible that bat faecal associated retrovirus 1 G04/aus/1 and bat faecal associated retrovirus 2 AdCPG/aus/1 constitute the first exogenous and intact betaretroviruses sampled from the faeces of bats in Australia. Unfortunately, virus identification through metatranscriptomics does not provide reliable information on whether a virus is endogenous and defective, or still functional and exogenous (Hayward et al., 2013, Hayward and Tachedjian, 2021). That the retroviruses detected here have all the necessary genes to comprise a functional virus, with undisrupted ORFs, were not detected in every library, and are not present in the bat genome, at the very least suggests that they are only recently endogenized and currently unfixed in the bat population. Further work confirming the nature of the retroviruses detected here is warranted since bats are known to be major hosts for retroviruses (Cui et al., 2015) and their cross-species transmission across mammalian orders is commonplace (Hayward et al., 2013).

In addition to mammalian viruses, we detected virus sequences that are likely invertebrate-associated. Of particular interest were those from the *Orthomyxoviridae* and *Reoviridae* that span a wide variety of hosts including mammals and were at high abundance in some of the Centennial Park libraries. Notably, bat faecal associated reovirus 1 CP02/aus/1 groups with members of the *Reoviridae* associated with ticks that are vectors for numerous pathogenic microorganisms, particularly from the genus *Coltivirus* – Colorado tick fever virus and Eyach virus (Goodpasture et al., 1978, Rehse-Küpper et al., 1976). Additionally, a novel coltivirus – Tai Forest reovirus – was sampled from bats in Cote d’Ivoire and shown to infect human cells (Weiss et al., 2017). The current evidence for tick-borne reovirus infection in humans highlights the importance of assessing the pathogenic potential of new tick associated reoviruses, especially those viruses discovered in urban wildlife.

Our study highlights the diversity of viruses in wildlife species from metropolitan areas. In this this context it is notable that the bat coronaviruses identified fall within the subgenus *Nobecovirus* of betacoronaviruses. Currently, this subgenus is strongly associated with bats sampled on multiple continents, with the phylogenetic depth of the *Nobecovirus* lineage further suggesting that bats have harbored these viruses for millennia with no apparent infection of humans. Hence, although the bats studied were resident in urban/suburban locations, this does not necessarily translate into a clear risk of human emergence.

## Data statement

The raw data generated for this study are available in the NCBI SRA database under the BioProject accession number PRJNA851532 and SRA accession numbers SRR19790899-SRR19790914. All genome sequences presented in phylogenetic trees are available in NCBI GenBank under the accession numbers ON872523-ON872588.

## Ethics statement

Ethics approval was granted by the University of Sydney Animal Ethics Committee (AEC 2018/1460).

## CRediT authorship contribution statement

**Kate Van Brussel**: Investigation, Data curation, Formal analysis, Writing – original draft, Writing – preparation, review and editing. **Jackie E. Mahar**: Formal analysis, Writing – preparation, review and editing. **Ayda Susana Ortiz-Baez**: Formal analysis, Writing – preparation, review and editing. **Maura Carrai**: Investigation, Writing – review. **Derek Spielman**: Investigation. **Wayne S. J Boardman**: Investigation, Writing – review. **Michelle L. Baker**: Resources, Writing – review. **Julia A. Beatty**: Investigation, Supervision. **Jemma L. Geoghegan**: Resources, Writing – review. **Vanessa R. Barrs**: Conceptualisation, Methodology, Funding acquisition, Supervision, Writing – review and editing. **Edward C. Holmes**: Conceptualisation, Funding acquisition, Supervision, Writing – preparation, review and editing.

