## Supplementary Information for "Faecal virome of the Australian grey-headed flying fox from urban/suburban environments contains novel coronaviruses, retroviruses and sapoviruses"

### Appendix A. Supplementary Information

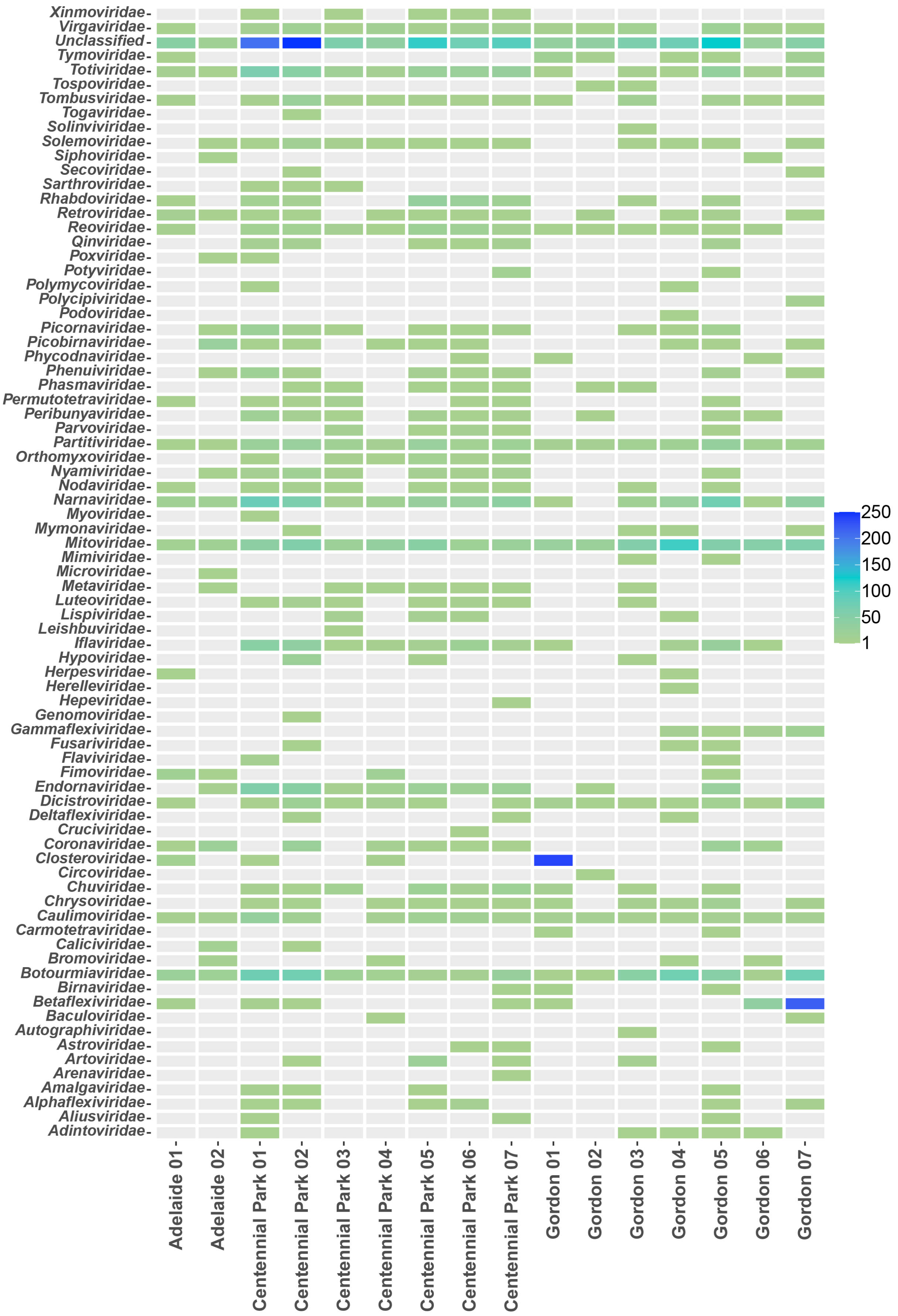

**Supplementary Fig. 1.** Overview of number of virus contigs for each sequencing library classified by virus family. The taxonomic classification was based on the taxonomy of the closest relatives in the NCBI non-redundant protein database.

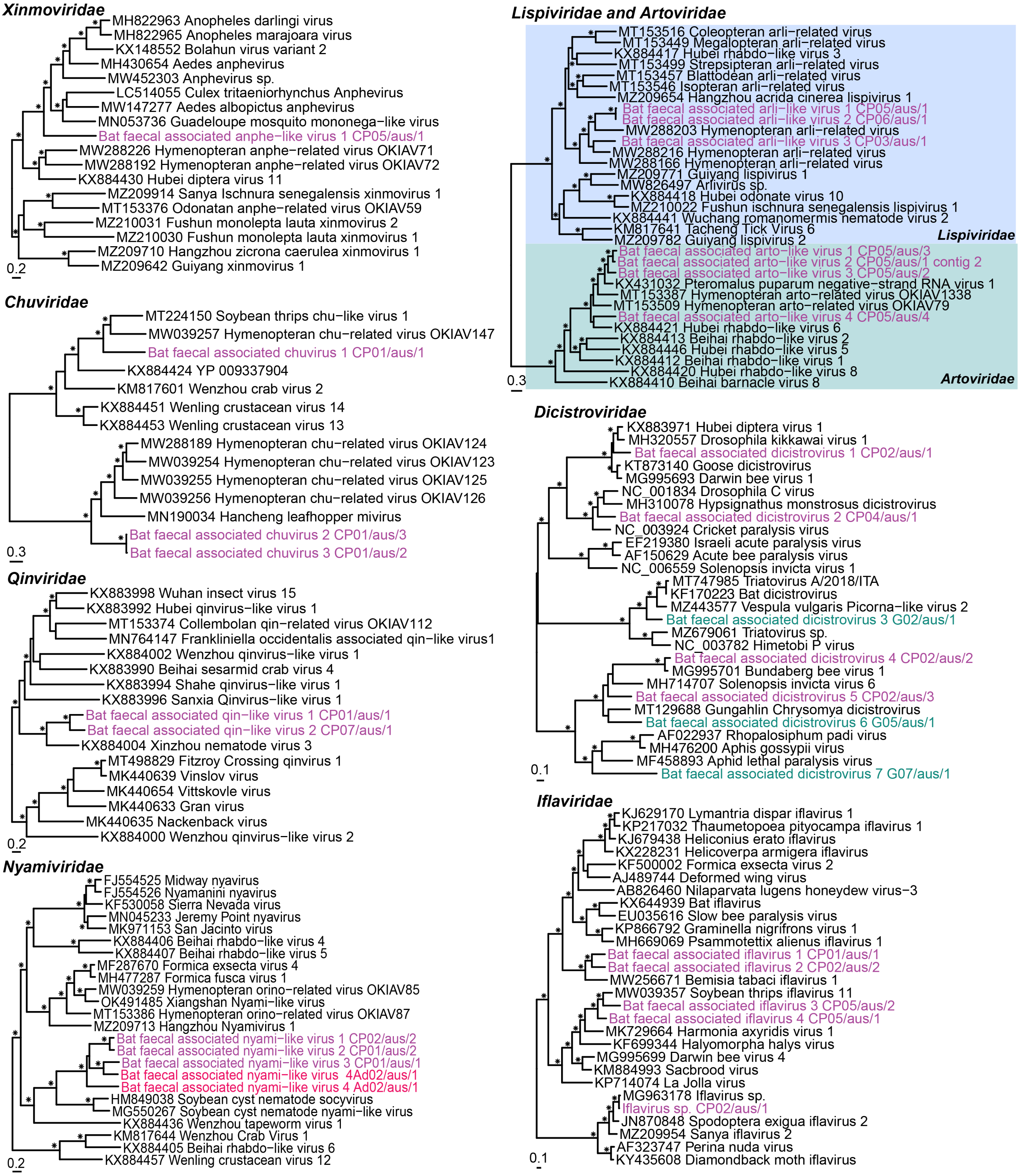


**Supplementary Fig. 2.** Phylogenetic analysis of the *Lispiviridae and Artoviridae*, *Nyamiviridae*, *Chuviridae*, *Xinmoviridae, Qinviridae, Iflaviridae* and *Dicistroviridae*. The RdRp amino sequence was used to estimate phylogenetic trees in all cases (alignment lengths of 1,778, 1,575, 2,066, 1,804, 1,179, 491 and 454 amino acid residues, respectively). Viruses from this study are coloured by sampling location (Adelaide – pink, Centennial Park – purple and Gordon – green), with the *Lispiviridae* and *Artoviridade* highlighted. Bootstrap values >70% are represented by the symbol shown at the branch node and the tree is midpoint rooted for clarity and the scale bar represents the amino acid substitutions per site.

**Supplementary Table 1.** VP1 amino acid identity of the novel sapoviruses in comparison to other bat sapoviruses.

| **Virus** | **Accession** | **Genogroup** | Amino acid identity (%) | |
|---|---|---|---|---|
| | | | Ad02/aus/4 | Ad02/aus/1 |
| Ad02/aus/4 | | GXX | NA | 77.7 |
| Ad02/aus/1 | | GXX | 77.7 | NA |
| Bat-SaV/Limbe65/CAM/2014 | KX759620 | GXIX | 29.6 | 38.8 |
| Bat-SaV/Limbe899a/CAM/2014 | KX759621 | GXIX | 30.7 | 39.9 |
| Bat-SaV/Limbe25/CAM/2014 | KX759618 | GXIX | 30.7 | 39.7 |
| Bat-SaV/Lysoka36/CAM/2014 | KX759619 | GXVIII | 19.8 | 32.1 |
| Bat-SaV/Limbe900/CAM/2014 | KX759623 | GXVIII | 29.8 | 37.5 |
| BtMm-CalV/JX2010 | KJ641703 | GXVII | 28.6 | 40.1 |
| BtCV/OV-157/M.dau/DK/2018 | MZ218056 | GXVII | 28.6 | 39.8 |
| BtCalV/M63/HUN/2013 | KU712497 | GXVII | 28.6 | 39.8 |
| BtRs-CalV-1/GX2012 | KJ641701 | GXVI | 25.8 | 36.5 |
| TLC39/HK | JN899074 | GXIV | 29.1 | 38.8 |
| TLC58/HK | JN899075 | GXIV | 29.4 | 38.6 |
| TLC34/HK | JN899072 | GXIV | 28.7 | 38.5 |
